# The role of elections as drivers of tropical deforestation

**DOI:** 10.1101/2021.05.04.442551

**Authors:** Joeri Morpurgo, W. Daniel Kissling, Peter Tyrrell, Pablo J. Negret, James R. Allan

## Abstract

Tropical forests support immense biodiversity and provide essential ecosystem services for billions of people. Despite this value, tropical deforestation continues at a high rate. Emerging evidence suggests that elections can play an important role in shaping deforestation, for instance by incentivising politicians to allow increased utilisation of tropical forests in return for political support and votes. Nevertheless, the role of elections as a driver of deforestation has not been comprehensively tested at broad geographic scales. Here, we created an annual database from 2001 to 2018 on political elections and forest loss for 55 tropical nations and modelled the effect of elections on deforestation. In total, 1.5 million km^2^ of forest was lost during this time period, and the rate of deforestation increased in 37 (67%) of the analysed countries. Deforestation was significantly lower in years with presidential or lower chamber elections compared to non-election years, which is in contrast to previous local-scale studies. Moreover, deforestation was significantly higher in presidential or lower chamber elections that are competitive (i.e. when the opposition can participate in elections and has a legitimate chance to gain governmental power) compared to uncompetitive elections. Our results document a pervasive loss of tropical forests and suggest that competitive elections are potential drivers of deforestation. We recommned that organisations monitoring election transparency and fairness should also monitor environmental impacts such as forest loss, habitat destruction and resource exploitation. This would benefit the tracking of potential illegal vote buying with natural resources.

## 1. Introduction

Tropical forests contain Earth’s richest biota and are the last refuges for many imperilled species (Gaston, 2000; Gibson et al., 2011). Tropical forests also provide globally important ecosystem services such as carbon sequestration and clean water provisioning (Foley et al., 2007). As many as 1.6 billion rural people live in close proximity to forests and may depend on forest resources for their livelihoods (Angelsen et al., 2014; Joshi and Joshi, 2019; Rudow et al., 2013). It is therefore concerning that tropical deforestation has reached critically high levels in the last few decades, with as much as 79,000 km^2^ – an area similar in size to Austria – being cleared every year (Austin et al., 2017). Understanding what drives tropical deforestation is thus crucial for implementing policy and conservation actions to ensure forest preservation.

The most prevalent direct causes of tropical deforestation include commercial logging (Curtis et al., 2018; Hosonuma et al., 2012), subsistence logging (e.g. for firewood; Heltberg et al., 2000; Hosonuma et al., 2012), conversion of forests to agricultural lands (e.g. for oil palm plantations or cropping; Hosonuma et al., 2012; Koh and Wilcove, 2008; Laurance et al., 2014), and wildfires which are often started by subsistence slash and burn agriculture (Laurance et al., 2002). There is good evidence that these drivers of deforestation increase when certain enabling factors are at play. One of these factors is corruption, which has been associated with higher rates of deforestation (Burgess et al., 2012; Smith et al., 2003; Wright et al., 2007). Another factor is the Gross Domestic Product (GDP) of a country, with higher deforestation occurring in countries with lower GDP (Ewers, 2006). Deforestation also tends to be higher in countries with higher human population densities (Sandker et al., 2017). Interestingly, factors such as a free media are associated with less deforestation, perhaps countering the effects of corruption (Bertot et al., 2010; Kolstad and Wiig, 2009). Other factors that potentially influence deforestation (e.g. armed conflicts, illegal crop production, or political elections and election cycles) have been less studied, even though there is growing evidence that they could drive deforestation trends in the tropics (Dávalos et al., 2016; Landholm et al., 2019; Negret et al., 2019).

Recent evidence suggests that elections could be key drivers of deforestation (List and Sturm, 2006; Pailler, 2018; Rodrigues-Filho et al., 2015). For example, a local scale study in Brazil found that municipal level deforestation was 8–10% higher in years when there was a municipal election (Pailler, 2018). Moreover, a similar increase in deforestation was also found during the national elections in Brazil (Rodrigues-Filho et al., 2015). During gubernational elections, in the United States of America, governors are more likely to advance or retract environmental policy based on the preference of the voters of their state. For instance, in “green” states environmental policy is more likely to advance during the election period, whereas in “brown” states it is more likely to retract (List and Sturm, 2006). A recent study investigating the economic and political incentives of deforestation in Indonesia found that deforestation substantially increases before a mayoral election, suggesting that political incentives reinforce tropical deforestation (Cisneros et al., 2021). This suggests that elections can influence deforestation, but broad generalizations should be made cautiously given the limited geographical scope, or the limited quality and resolution of deforestation data used in these studies so far.

Elections could increase deforestation via multiple mechanisms. Elections are power struggles where politicians aim to gain an advantage over opponents. These advantages can be achieved through popular policies and by creating economic opportunities (Akhmedov and Zhuravskaya, 2003; Drazen and Eslava, 2010; Nordhaus, 1975). For example, politicians might gift or promise forested land for exploitation to win favour with powerful potential supporters, or with businesses such as developers and loggers. A real world example occurred in Uganda in 2011, where the incumbent government promised forests to win community support (Médard and Golaz, 2013). A similar example is the 2018 Brazilian presidential elections which caused a spike in deforestation due to candidates promising the dismantling of environmental laws (Abessa et al., 2019). Leading up to elections, governments may be so focussed on electioneering that diverts their attention from environmental protection and turn a blind-eye to people utilising forest resources, allowing them to harvest unsustainably or to settle on protected forested land (Negret et al., 2017). Most countries have strong laws against winning political favour through financial bribery. However, environmental protection laws are usually less rigorously monitored or upheld than financial laws, making winning support by giving away land and forest resources an attractive alternative to money (Ohman, 2013). There are many mechanisms for elections to drive deforestation but the effect of elections on deforestation remains under-investigated, especially at broad geographic extents.

Here, we analyse the effect of elections as drivers of deforestation at a pantropical scale. We focus on the tropics because the mechanisms and drivers of deforestation are fairly distinct from the higher latitude forests in the temperate, boreal and taiga zone (Curtis et al., 2018). To assess the drivers of tropical deforestation, we first explored the directionality and shape of temporal trends in deforestation within 55 pantropical countries from 2001 to 2018 using remotely-sensed global forest loss data (Hansen et al., 2013). High-resolution (30 × 30 metre) year-by-year global forest loss data is now available from 2000 to 2018 (Hansen et al., 2013), providing new opportunities to study the effect of elections on deforestation more accurately and at unprecedented spatial extents.

We created an annual database over this time period covering the year in which national elections took place and which type of election it was (presidential, lower chamber, and upper chamber elections). We further extracted additional information on governance (e.g. competitiveness, media integrity, corruption control) and human population density. We used a Hierarchical generalized additive model (HGAM; Pedersen 2019) to assess the effect of election and the governance variables on the proportional deforestation of countries relative to their forest cover in the year 2000. This HGAM approach allows the modelling of non-linear functional relationships between covariates and outcomes where the shape of the function itself varies between different grouping levels (e.g. countries). This technique allowed us to disaggregate the changes in forest loss in each country over time - which can be driven by various factors - from the election covariates. These analyses allowed us to (1) quantify the effect of presidential, lower chamber, and upper chamber elections on tropical deforestation rates compared to non-election years, and (2) to test whether the competitiveness of an election has an effect on deforestation.

## 2. Methods

### 2.1 Data collection

We developed an annual 2001–2018 database for 55 tropical-forest countries (Table A1; Figure A1) covering national and state-level deforestation, election dates, governance variables and human population density. The governance variables included competitiveness of elections, media integrity of a country, and control of corruption (Table 1). Human population density captured the number of residents per country area (Table 1).

**Table 1.** Summary of predictor variables which were included in Hierarchical Generalized Additive Models to explain proportional deforestation of a country relative to the forest cover in the year 2000 (response variable). The predictor variables capture governance aspects (competitive elections, media integrity and control of corruption) and human population density.

| Variable | Definition and methods | Reference & source |
| --- | --- | --- |
| Competitive elections | Competitive elections (referred to as 'Competitive' in our analysis) is a binary variable that quantifies whether elections are sufficiently free for the opposition to gain legislative or executive power (1) or not (0). This reflects whether the seats of the executive and legislative body are filled by elections that are characterized by uncertainty in terms of the final outcome. This includes that (1) the legislature is only constitutionally dissolved, (2) members of the executive or legislative are only constitutionally removed, (3) elections are held at a time consistent with constitutional requirements, (4) non-extremist parties are not banned, and (5) voters experience little systematic coercion in their electoral vote. | Tufis, 2019 |
| Media integrity | Media integrity measures to what extent media are diverse and critical on governmental issues. It is a continuous composite variable with a range 0.00–0.83, based on five indicators: (1) How often media are critical of the government, (2) how wide the range of media perspectives is, (3) if there is media bias against government opposition, (4) whether media accepts bribes to alter news coverage, and (5) to what extent criticism of the government is common and normal in the mediated public sphere. | Tufis, 2019 |
| Control of corruption | Control of corruption measures the perception of corruption by public power for private gain. It is an continuous index with a range -1.68–0.76, created by modelling 50 variables on corruption. It intends to capture the extent to which public power is exercised for private gain. This includes both petty and grand forms of corruption, and the 'capture' of state assets by elite and private interests. | Kaufmann et al., 2011 |
| Human population density | Population density is defined as all residents of a given political unit divided by its area (i.e. individuals per km<sup>2</sup> of terrestrial land of a country). Refugees who are not permanently settled are excluded. The variable is continuous and ranges 3.05–498.66. | World bank, 2020 |

We extracted annual forest loss data for each country for the years 2001–2018 using high resolution (30 × 30 metre) global maps of forest cover and forest loss (Hansen et al., 2013). Data were extracted and processed in the Google Earth Engine (https://earthengine.google.com), a cloud platform for earth-observation data analysis (Gorelick et al., 2017). We adapted code from Tracewski et al. (2016) to quantify forest loss per year and country, and make our code available via GitHub (https://github.com/JoeriMorpurgo/Elections2020). The Global Forest Change database defines forest as >50% crown cover of trees taller than 5 m height. The presence of forest is given for each 30 × 30 metre pixel using the year 2000 as a baseline. Forest loss is defined as the disappearance of a forest pixel in a given year (1 = loss, 0 = no loss). A given forest pixel can only be lost once (in years 2001–2018). We used the available data on forest cover (year 2000) and forest loss (years 2001–2018) to calculate the proportional loss (i.e. deforestation) over a given year within (sub)national boundaries relative to the forest cover in the year 2000 (see methodological example in Figure A1). We did not include ‘gain’ in forest area because it is only provided as a total over the whole time period (Hansen et al., 2013) and because it is often due to plantation forests rather than natural regrowth or restoration (Tropek et al., 2014). The Global Forest Change data is considered the most accurate global deforestation data available. However, we acknowledge limitations such as the inability to differentiate between forest and agro-forests, which have been discussed elsewhere (Tropek et al., 2014, Allan et al. 2017).

We gathered data on when national level elections took place by examining each country’s constitution, and cross-checking this with a number of election databases (see Table B2). In the few cases where we could not find a formal source we utilised Wikipedia (n = 4, 0.9%), which is regarded as a credible source for election data (Brown, 2011). We collected information on three types of national elections: (i) *Lower chamber elections*, where the lower chamber holds the legislative power allowing them to create laws; (ii) *Upper chamber elections*, where the upper chamber reviews the legislative power; and (iii) *Head-of-state or head-of-government elections* (hereafter called *‘presidential elections’*) depending on who holds the executive power to enforce the law and is elected. All countries analysed had a lower chamber and presidential elections. However, many countries did not have upper chamber elections (25 out of 55 countries, i.e. 45%). Presidential and upper chamber election dates often occur in the same year as lower chamber elections (52% and 38% of the time, respectively). All election types were treated as a binary predictor variable (1 = year with election, 0 = no election), i.e. either occurring in a given year or not.

We extracted governance information and human population density from various sources (for details see Table 1). Elections were scored as competitive (= 1) when they are sufficiently free for the opposition to gain legislative or executive power with enough votes, and otherwise as non-competitive (= 0) (see ‘Competitive elections’ in Table 1). Note that this variable does not capture whether parties have equal funding, media coverage or whether civil liberties are respected. Hence, competitive elections are not equal to free and fair elections (Skaaning et al., 2015). We further used an index from the World Bank which captures the control of corruption, which has been linked to both tropical deforestation and enhancing election cycles (Kaufmann et al., 2011; Pereira et al., 2009; Smith et al., 2003). We also extracted a variable which quantifies to what extent media are diverse and critical (‘Media integrity’ in Table 1), as this has been shown to counter the effects of election cycles (Akhmedov and Zhuravskaya, 2003; Tufis, 2019). Finally, we also accounted for human population density, since higher densities at a national level tend to increase deforestation (World bank, 2020). All predictor variables, included in the analysis, were compiled at national and annual scale. Four countries lacked data on ‘Competitive elections’, leading to exclusion in the Hierarchical Generalized Additive Modelling (Table A1).

### 2.2 Statistical analyses

The statistical analysis aimed to assess (1) the directionality and shape of temporal trends in deforestation, (2) the effect of presidential, lower chamber, and upper chamber elections on deforestation, and (3) the effect of competitiveness of elections on deforestation trends.

First, we used a non-parametric Mann-Kendall test (Kendall, 1938; Mann, 1945) to test for monotonic trends (i.e. directionality) of deforestation over time for each country. This test is more robust to outliers, non-normality and temporally autocorrelated data than simple linear models and is widely used in time-series analysis (Yue et al., 2002).

Second, we used Hierarchical Generalized Additive Models (HGAM) (Lin and Zhang, 1999; Pedersen et al., 2019; Wood, 2017) to model non-linear trends in deforestation in relation to election type and competitiveness of elections. The flexible nature of HGAMs allows for modelling smooth patterns across space and over time, with the amount of smoothing controlled to prevent over-fitting (Wood 2017). The HGAM approach thus allows the modelling of non-linear functional relationships between covariates and outcomes where the shape of the function itself varies between different grouping levels. In our case, this grouping variable was the country level. This technique allowed us to disaggregate the changes in forest loss in each country over time - which can be driven by various factors - from the election covariates. Our models used a global smoother plus country-level smoothers with differing wiggliness (Pedersen et al., 2019).

We used three separate HGAMs to model each election type independently: a presidential model, a lower chamber model and an upper chamber model. The general mathematical formulation of the HGAMs was:

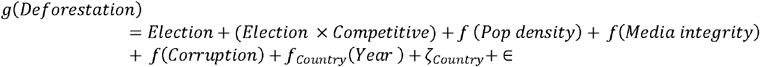

Where *g*(*Deforestation*) is the response variable defined as proportional deforestation of a country relative to the forest cover in the year 2000. The binary predictor variable *Election* is 1 when an election is being held in a given year, and 0 if not. The *Election* term differs among HGAMs because of the different election data (presidential, lower chamber or upper chamber). The binary predictor variable *Competitive* is 1 if the election setting is competitive, and 0 if it is not, and modelled as an interaction with the *Election* term (i.e. *Election* × *Competitive*). The predictors *f* (*Pop density_i_*), *f* (*Media integrity_i_*) and *f* (*Corruption_i_*) are all modelled smooths allowing for non-linear relationships. All smooths used penalized thin plate regression splines (TPRS) (Wood, 2003). With these splines, the null space is also penalized slightly, and the whole term can therefore be shrunk to zero, effectively acting as a model fitting step (Wood, 2003). The additional advantage of the TPRS approach is that knot positions were selected automatically from the data, eliminating knot placement subjectivity. Random effects are described by *ζ_Country_*, which accounts for country-level mean differences of deforestation at the intercept as suggested by Pedersen et al. (2019). The term *f_Country_* (*Year*) is a separate univariate smooth for each country to account for intergroup variability. We used a Gaussian process smooth to account for temporal autocorrelation (Wood, 2017). Finally, ∈ describes the error that is not explained by the other terms. HGAMs were modelled using a beta regression logit link structure to account for the proportional nature of the response variable which is bound between 0 and 1, and overcomes limitations in other more commonly used distributions (Douma and Weedon, 2019). For each term the penalty controlling the degree of smoothing was selected using restricted maximum likelihood (REML; Wood 2017, p. 185)

The autocorrelation function of the residuals, concurvity and model residuals were visually inspected for all models, and no issues were identified. The supplementary material provides the autocorrelation function of the residuals (Figure B1), the concurvity (Figure C 1–3), and the model residuals (Figure D1).

## 3. Results

### 3.1 Global deforestation trends from 2001 to 2018

We found that 1.5 million km^2^ of tropical forest – an area similar in size to Mongolia – was lost between 2001 and 2018 in the 55 tropical countries analysed (Table 1A). The largest area of forest loss occurred in Brazil (469.839 km^2^), followed by Indonesia (227.008 km^2^) and the Democratic Republic of Congo (112.626 km^2^) (Figure 1A). On average, 0.52%of the world’s tropical forests were lost each year from 2001 to 2018 (SD = 0.15%, range = 0.35%–0.91%, *n* = 55 countries). The overall proportion of pantropical deforestation has increased during this time by 182%, with 37 out of the 55 assessed countries (67%) showing statistically significant increases (demonstrated by Mann Kendall tests showing statistically significant positive tau values at *p* < 0.05) (Figure 1B). Only four countries decreased in their annual rate of deforestation (indicated by negative tau values of the Mann Kendall tests), but these were statistically not significant (at *p* > 0.05).

**Figure 1.**
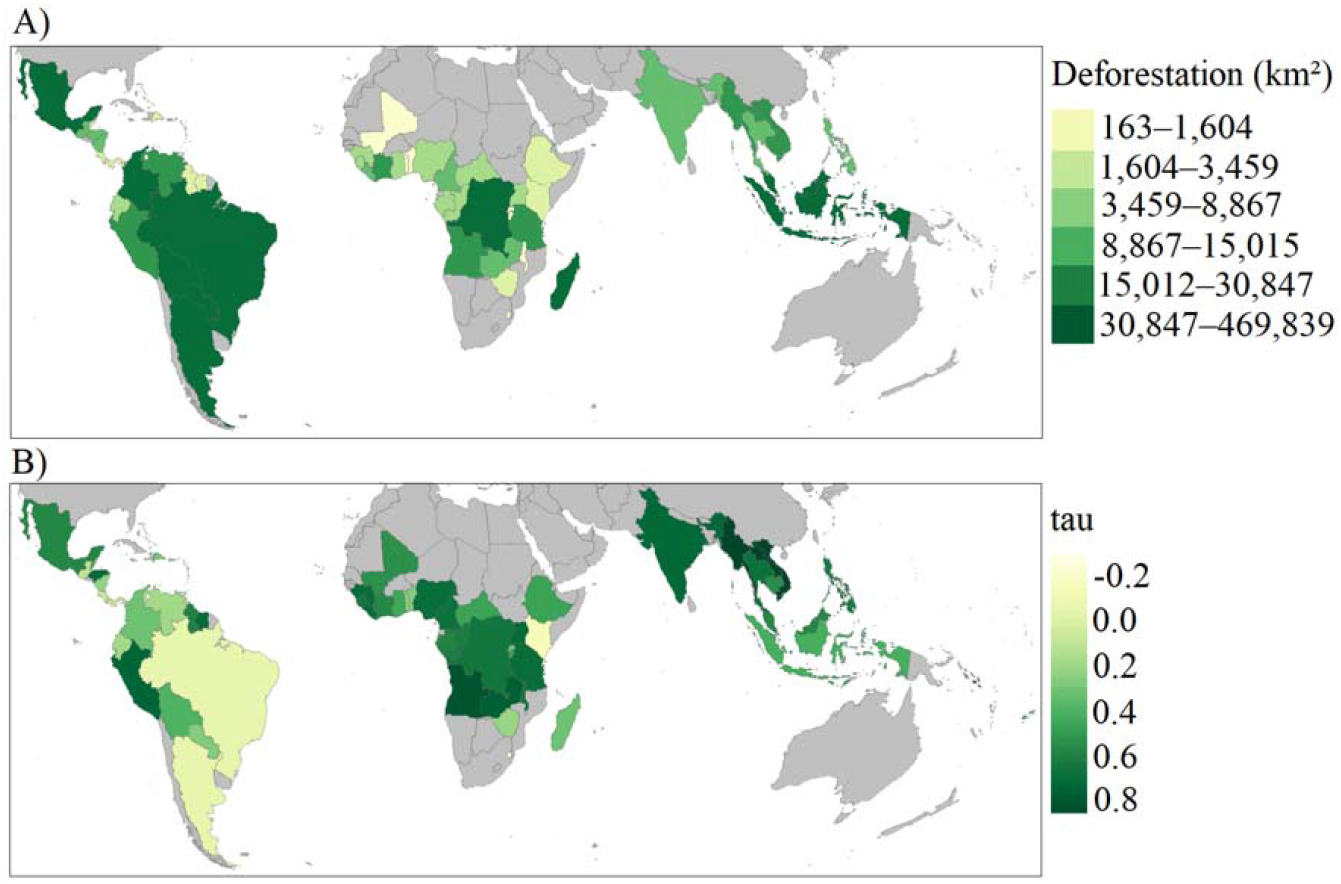
Deforestation in 55 tropical countries between 2001–2018. A) Total amount of deforestation (in km^2^) at a national scale from 2001–2018. B) Directionality and strength of national deforestation trends quantified as correlation coefficients (Tau values) from Mann Kendall tests. A total of 51 countries show an increase in the annual rate of deforestation (light green-green: positive Tau values) whereas four countries show a decrease (light yellow: negative Tau values). Annual forest loss data for each country were derived from high resolution (30 × 30 metre) global maps of forest cover and forest loss (Hansen et al., 2013).

The shapes of deforestation trends derived from the HGAMs varied considerably among countries (*n* = 51) (Figure 2A). In general, they followed five main typologies (Figure 2B–F): linearly increasing, linearly decreasing, curvilinearly increasing, curvilinearly decreasing and fluctuating. We visually inspected these deforestation trends for each country and found that the rate of deforestation increased in 36 (71%) of the analysed countries (*n* = 51). Of those, 24 countries showed a linearly increasing deforestation trend (Figure 2B) and 12 countries an increasing curvilinear trend (Figure 2D). Two countries showed a linearly decreasing trend (Figure 2C) and five countries curvilinearly decreasing trend (Figure 2E). A total of 8 countries were classified as having fluctuating deforestation trends (Figure 2F).

**Figure 2.**
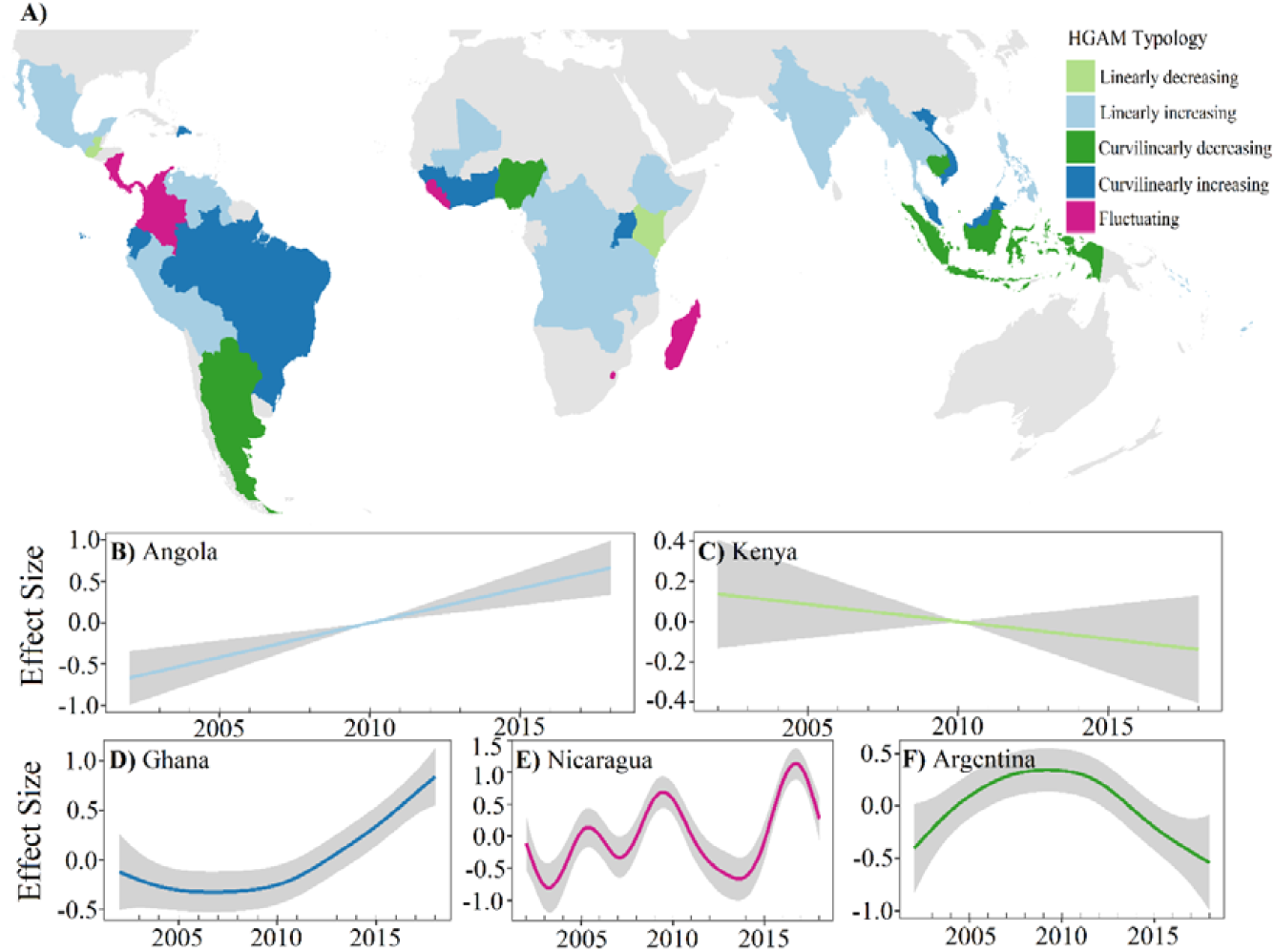
National deforestation trends between 2001–2018 across the tropics. A) Pantropical overview of main typologies of deforestation trends (linearly increasing, linearly decreasing, curvilinearly increasing, curvilinearly decreasing and fluctuating) as derived from Hierarchical Generalized Additive Models (HGAMs). Examples of trend typologies: B) linearly increasing (Angola), C) linearly decreasing (Kenya), D) curvilinearly increasing (Ghana), E) fluctuating (Nicaragua), and F) curvilinearly decreasing (Argentina).

### 3.2 Election types and deforestation

All three HGAMs had high explanatory power (R^2^ > 0.87, explained deviance > 90%, see Table 2) and show that tropical deforestation is lower in years when there is a presidential or lower chamber election, compared to years with no election (Figure 3A, B). This is demonstrated by the negative and statistically significant logit estimate for in the two HGAMs for presidential and lower chamber elections (Table 2). The logit estimate for the upper chamber HGAM also showed a negative sign but was statistically not significant (Table 2, Figure 3C).

**Figure 3.**
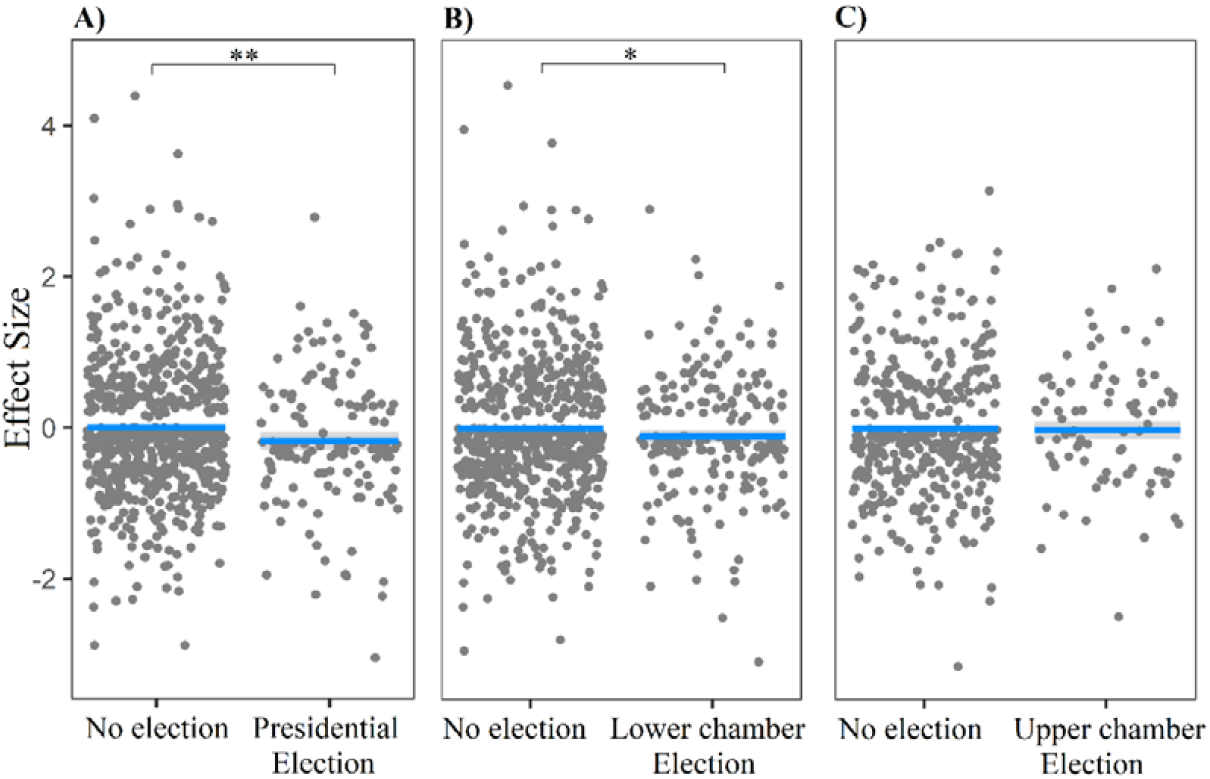
The effect of elections on deforestation. Each point represents the amount of logit-transformed deforestation in a given year and country. The presidential election and lower chamber election (A and B) show statistically significant lower deforestation compared to non-election years (compare results for term in Table 2). Upper chamber elections (C) show a similar but statistically not significant trend.

**Table 2.** Results of Hierarchical Generalized Additive Models (HGAMs) with a logit-link to explain the proportional deforestation of a country relative to the forest cover in the year 2000 (response variable). Three different HGAMs were implemented depending on the specific election type (presidential, lower chamber, or upper chamber election). Binary predictor variables are shown with parametric coefficients (logit estimates) whereas continuous variables are represented with smooth terms. For details of predictor variables see Table 1. Country-level estimates (n = 51 countries) were excluded from this table. Statistically significant p-values (p < 0.05) are indicated in bold.

|  | Presidential model |  |  |  | Lower chamber model |  |  |  | Upper chamber model |  |  |  |
| --- | --- | --- | --- | --- | --- | --- | --- | --- | --- | --- | --- | --- |
| Predictor | *Estimate* | *Std. error* | *Z-value* | *p* | *Estimate* | *Std. error* | *Z-value* | *p* | *Estimate* | *Std. error* | *Z-value* | *p* |
| *Intercept* | -5.47 | 0.10 | -57.10 | **<0.001** | -5.47 | 0.10 | -56.90 | **<0.001** | -5.52 | 0.14 | -40.34 | **<0.001** |
| Parametric coefficients |  |  |  |  |  |  |  |  |  |  |  |  |
| *Election* | -0.19 | 0.06 | -3.07 | **0.002** | -0.10 | 0.04 | -2.33 | **0.02** | -0.02 | 0.06 | -0.35 | 0.73 |
| *Election × Competitive* | 0.16 | 0.07 | 2.30 | **0.02** | 0.13 | 0.06 | 2.26 | **0.02** | -0.01 | 0.07 | 0.09 | 0.93 |
| Smooth term |  | *edf* | *Chi<sup>2</sup>* | *p* |  | *edf* | *Chi<sup>2</sup>* | *p* |  | *edf* | *Chi<sup>2</sup>* | *p* |
| *f (Population density)* |  | <0.001 | <0.001 | 0.79 |  | <0.001 | <0.001 | 0.82 |  | <0.001 | <0.001 | 0.36 |
| *f (Media integrity)* |  | <0.001 | <0.001 | 0.41 |  | <0.001 | <0.001 | 0.37 |  | 4.70 | 454.09 | 0.39 |
| *f (Control of corruption)* |  | <0.001 | <0.001 | 0.70 |  | <0.001 | <0.001 | 0.67 |  | <0.001 | <0.001 | 0.70 |
| Adjusted R<sup>2</sup> | 0.87 |  |  |  | 0.87 |  |  |  | 0.91 |  |  |  |
| Explained deviance | 90.2% |  |  |  | 90.1% |  |  |  | 93.2% |  |  |  |

### 3.3 Effect of competitiveness on deforestation

Deforestation was significantly higher in competitive presidential and lower chamber election years, compared to non-competitive election years (Figure 4A, B). This is demonstrated in the positive and statistically significant interaction term *Election* × *Competitive* in the presidential and lower chamber HGAMs (Table 2). The upper chamber HGAM showed a negative interaction term *Election* × *Competitive* but this was not statistically significant (Table 2, Figure 4C). None of the other predictors (human population density, media integrity, and control of corruption) showed a statistically significant effect on deforestation trends (Table 2).

**Figure 4.**
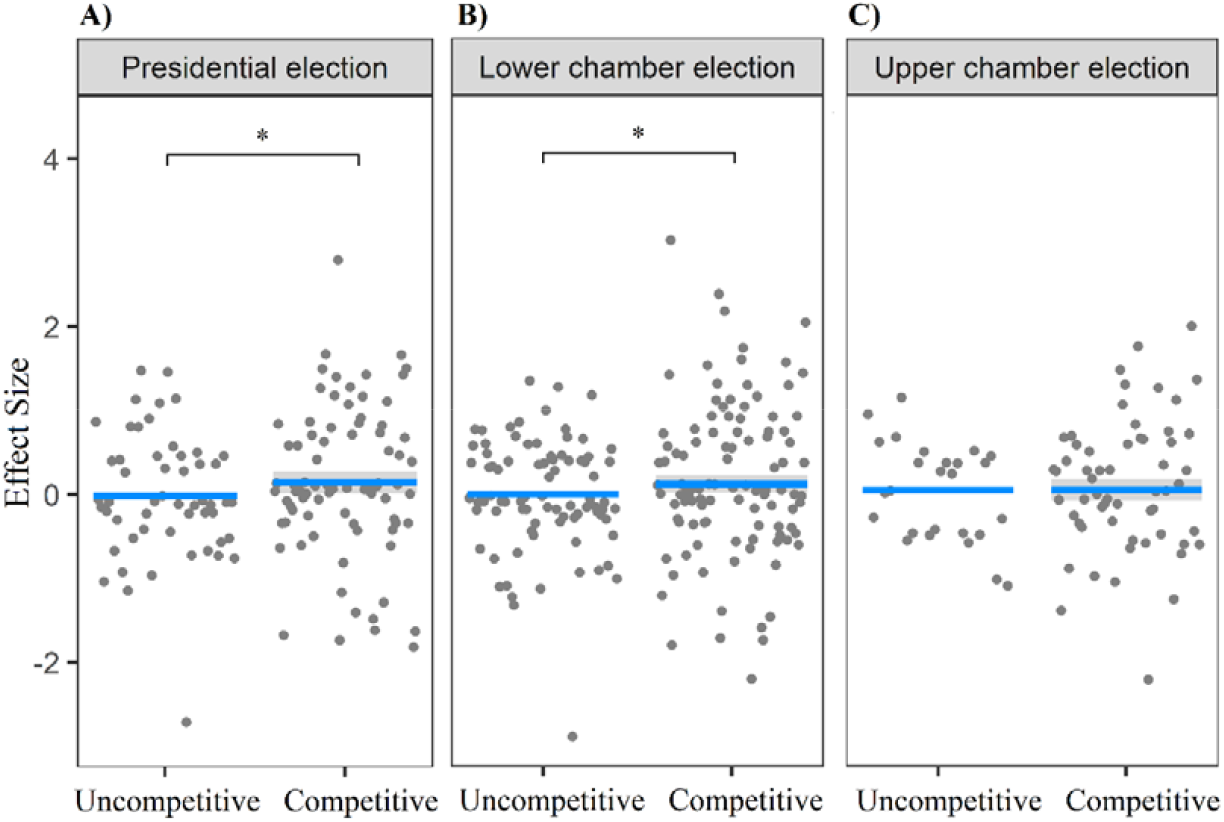
The effect of competitive elections on deforestation. Each point represents the amount of logit-transformed deforestation in an election year by country. Competitive presidential and lower chamber elections (A, B) show statistically significant higher deforestation compared to uncompetitive elections (compare results for interaction term in Table 2). Upper chamber elections (C) show an opposite but statistically not significant trend.

## 4. Discussion

We analysed the effect of elections on deforestation in 55 tropical countries over an 18-year time period and found that competitive elections are potential drivers of pantropical deforestation, whereas a general effect of election years on deforestation was not confirmed. We focused on tropical forests because they are predominantly lost due to agriculture and commodities exploitation rather than changes caused by forestry management and wildfires, as for instance in boreal and taiga forests (Curtis et al., 2018). Our results document a massive tropical forest loss from 2000 to 2018, with the rate of deforestation increasing in more than two-thirds of the studied countries. To our knowledge, this represents the most comprehensive and most up-to-date study of its kind covering a pantropical extent.

Our results confirm the well-known trend that deforestation rates are increasing across the tropics (Curtis et al., 2018). This is alarming since deforestation is accelerating while the remaining forest area is becoming smaller. While the majority of studied countries (70.5%, HGAM) show linearly or curvilinearly increasing deforestation trends, there are a few countries in which deforestation is decreasing (13.7%) or fluctuating with sporadic increases and decreases (15.6%). The decreases observed in our results seem to coincide with the implementation of forest protection policies or actions. For example, in 2004, The Brazilian Government founded the country’s environmental enforcement agency (IBAMA) which led to a 37% reduction in deforestation between 2005 and 2007, by using in field enforcement using real time deforestation detection (Arima et al., 2014; Soares-Filho et al., 2010). Similarly, conservation in the Colombian Guyana Shield reduced deforestation in protected area’s compared to their buffer zone, with only 1% of the natural forest in protected areas lost, while the buffer zones lost between 5–7% between 1985 and 2002. In particular, the reduced deforestation in protected areas was linked with lower infrastructure, accessibility and reduction in illicit agriculture (Armenteras et al., 2009). These examples are encouraging since they show that policy tools and conservation intervention can effectively limit deforestation and that governments have the means to take the necessary steps to halt ongoing deforestation (Busch et al., 2015; Rudorff et al., 2011; Umemiya et al., 2010; Wehkamp et al., 2018).

In contrast to our expectation, we found that deforestation was significantly lower in election years than in non-election years. This is, for instance, opposite to a study at the national scale of Brazil where municipal-level deforestation increased by 8–10% in years with a municipal election (Pailler 2018). In general, election theory suggests that politicians should utilise all avenues possible to win support and favour in the lead up to an election, which includes giving away or promising forested land for development, or turning a blind eye to forest exploitation (Abessa et al., 2019; Akhmedov and Zhuravskaya, 2003; Burgess et al., 2012; Shi and Svensson, 2006). We suggest that there are several plausible explanations why our analysis did not show such an expected effect.

First, forested land might be exploited before an election, causing an increase in deforestation above baseline levels. This exploitation may then stop shortly after an election, and result in a decrease of deforestation. This has been observed in some countries, for example in Russia, where election cycles in social expenditure from local governments generally drop one month after the election (Akhmedov and Zhuravskaya, 2003). This makes it difficult to detect a signal of elections on deforestation rates when analysing deforestation in yearly intervals since the pre-and post-election increase and decrease could cancel each other out. The global forest loss data that we used is currently only available in yearly intervals (Hansen et al., 2013) and thus does not account for short term pre/post-election changes in deforestation rates (e.g. within years). More detailed data (e.g. capturing intra-annual variation in deforestation rates during election periods) could benefit future studies of forest loss at national and global scales.

A second plausible reason for not detecting an increase of deforestation with presidential, lower and upper chamber elections is that forest governance and natural resource management is increasingly becoming decentralised within countries (Ginsburg and Keene, 2020). In principle, decentralisation should make it more difficult for national level politicians to exploit locally managed resources (Busch and Amarjargal, 2020). It is difficult to account for this decentralisation in global analyses because appropriate data to quantify the degree of local or municipal autonomy in forest management are lacking. Additionally, election data on the subnational administrative units that manage the forests are often missing. Hence, it is possible that local governments protect their forests from exploitation by higher level politicians during national election years. An alternative way to study this could be to investigate the effect of elections on deforestation at the spatial scale at which forest management decision are made. For instance, if forests are managed at the state or county level, the effects of state or county-level elections on deforestation could be analysed. To our knowledge, there are currently no global databases available that specify the level and spatial scale of forest governance.

Besides the general effect of presidential, lower and upper chamber elections on deforestation, our analysis revealed that deforestation is significantly higher in competitive election years compared to non-competitive election years. This supports our expectation that more competitive elections will increase incentives for politicians to misuse public goods for winning favour (Sanford, 2019; Shi and Svensson, 2006). To improve forest protection, we recommend that integrity and transparency monitoring schemes for elections such as the Global Network of Domestic Election Monitors (GNDEM) extend their mandate to include monitoring natural resources such as forests (Pereira et al., 2009; Shi and Svensson, 2006). Conservation groups should also remain vigilant during the lead up to elections, especially given land gifting practices for forest exploitation during the elections of Uganda in 2011 (Médard and Golaz, 2013).

## 5. Conclusions

Protecting biodiversity in tropical forests and their ecosystem services is crucial for meeting international policy targets such as the United Nations Sustainable Development Goals (SDGs) and the post-2020 targets of the Convention on Biological Diversity (CBD). Our analysis shows that tropical forests continue to decline and that elections can at least partly play a role in driving deforestation trends. However, more detailed data on intra-annual variation of deforestation and the spatial scale of forest governance are needed to improve our global (pantropical and cross-national) understanding of how elections influence forest loss driven. We urge electoral management bodies and conservation groups to be vigilant during competitive elections, because forests and other natural resources could be traded for votes. Further elucidating the role of elections on deforestation should be a focus of forest conservation efforts.

## Supporting information

Figure A1. Proportional deforestation

Figure B1. ACF

Figure C1-3. Concurvity

Figure D1. Residuals

Table A1. Countries

## 6. Data accessibility

Data and code used for this study have been made permanently and publicly available on the Mendeley Data repository at http://dx.doi.org/10.17632/5ngc9n3shd.1.

