## Supplementary material for "The role of elections as drivers of tropical deforestation": Figure A1. Proportional deforestation

Supplementary text 1. Methods of extracting forest loss for countries that changed subnational boundaries.

Between 2001 and 2018, twenty countries changed their state borders. To accurately assess state-level deforestation we used multiple maps, including ones from before and after the border changes (Retrieved from; Natural Earth, 2018; GDAM, 2018; Humanitarian Data Exchange, 2018, 2019; IPUMS International, 2012; FEWS, 2009, 2016). For example, when a country changed its state borders in 2008, we did our forest loss calculation using a map of the borders from 2001 to 2007, and then post 2007, used a different map of the state borders from 2008 to 2018 (Figure 5). We therefore calculated the proportion of forest-loss in a given year relative to the forest coverage of the year 2000, from the corresponding map. This method allows us to join the data from the pre- and post-border change maps accurately.

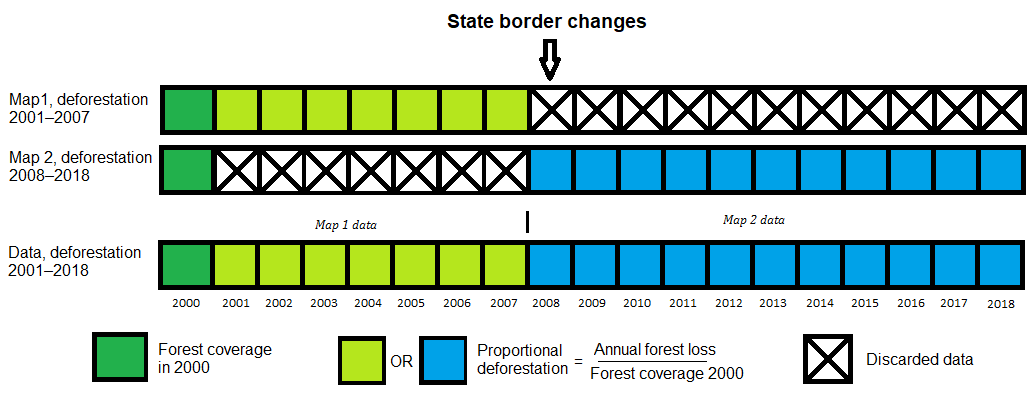

**Figure A1. Example of calculating the proportional deforestation for a country with state-level border changes.** The upper two bars represent the maps used to calculate the proportional deforestation covering 2001–2007 and 2008–2018. The final bar represents the data used in the analysis. National level deforestation, using these maps, was calculated by dividing the forest loss in a given year by the forest coverage in year 2000 of its corresponding map. In this example, data on forest loss used in the analysis would be two-part where 2001–2007 forest loss originates from map 1, and is joined with the 2008–2018 forest loss which originates from map 2.
