## Supplementary material for "The role of elections as drivers of tropical deforestation": Figure B1. ACF

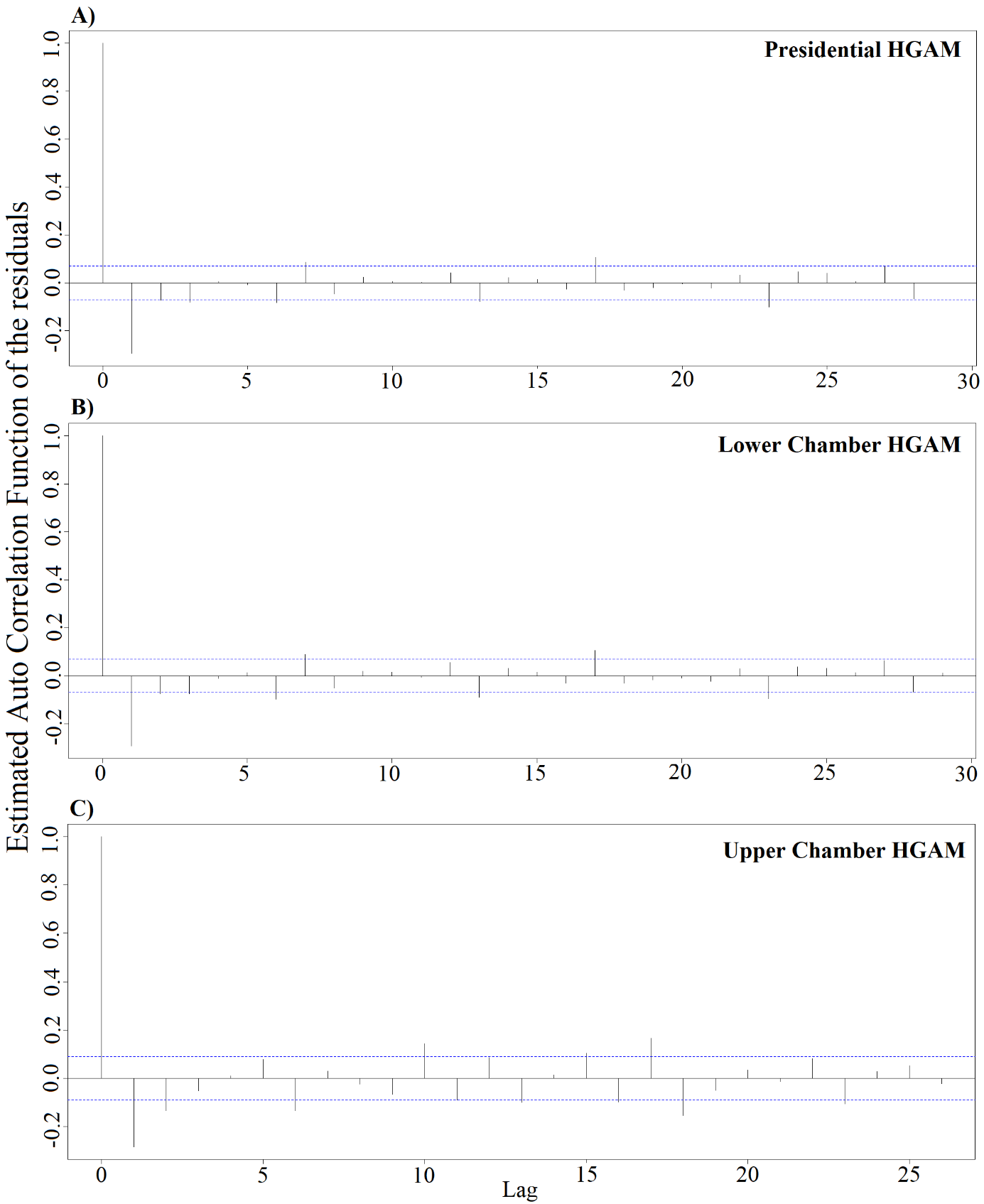

**Figure B1. Shown is the estimate Auto Correlation Function (ACF) of the residuals from the HGAMs.** The conventional 0.1 threshold is represented by the blue dotted line, indicating autocorrelation of the residuals. The models, except for the first lag value, remain mostly within this threshold indicating that there is no evidence of an autocorrelation impacting the models. This suggests that the models properly fit. The (A) Presidential, (B) Lower chamber and (C) Upper chamber strongly resemble each other, with the Upper chamber model being more volatile.
