## Supplementary material for "The role of elections as drivers of tropical deforestation": Figure C1-3. Concurvity

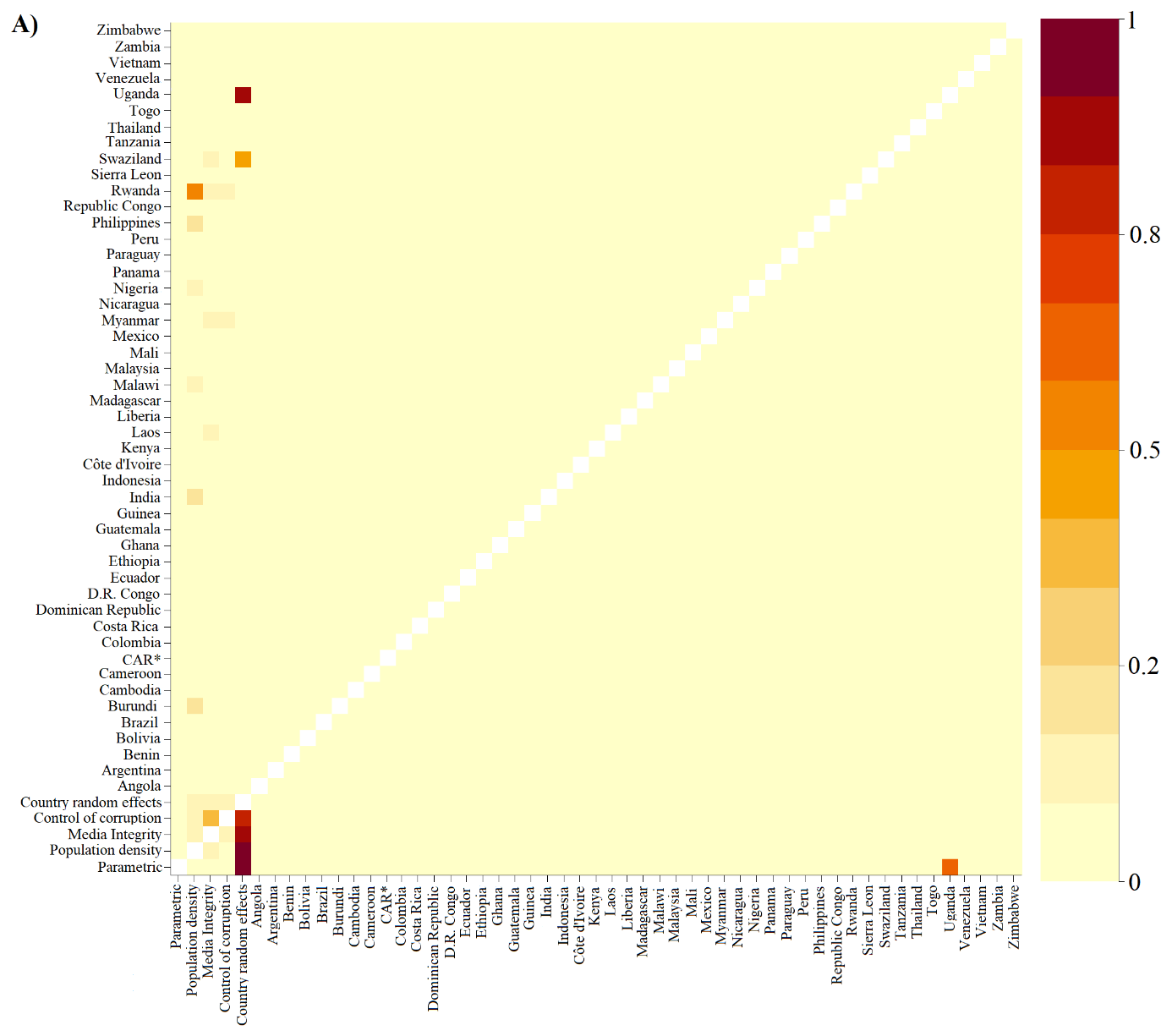

**Figure C1. Estimated concurvity from the presidential HGAM.** High values of concurvity were attained in relation to the country random effects term. The rest of the concurvity diagnostics have low values indicating that the model term do not explain each other well. This suggest that the model does not have problems associated with high concurvity, which are similar to high co-linearity. * - Central African Republic.

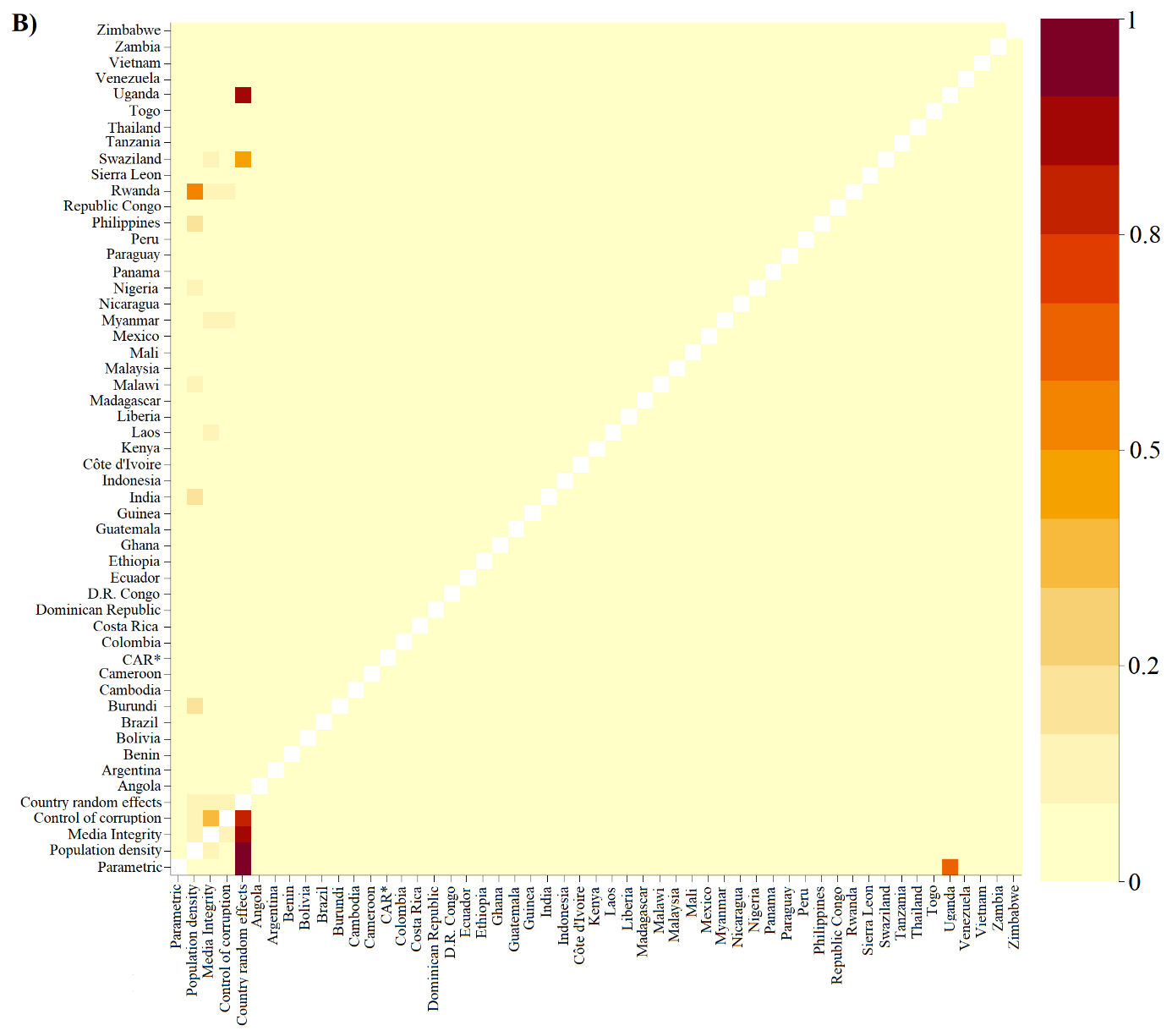

**Figure C2. Estimated concurvity from the lower chamber HGAM.** High values of concurvity were attained in relation to the country random effects term. The rest of the concurvity diagnostics have low values indicating that the model terms do not explain each other well. This suggest that the model does not have problems associated with high concurvity, which are similar to high co-linearity. * - Central African Republic.

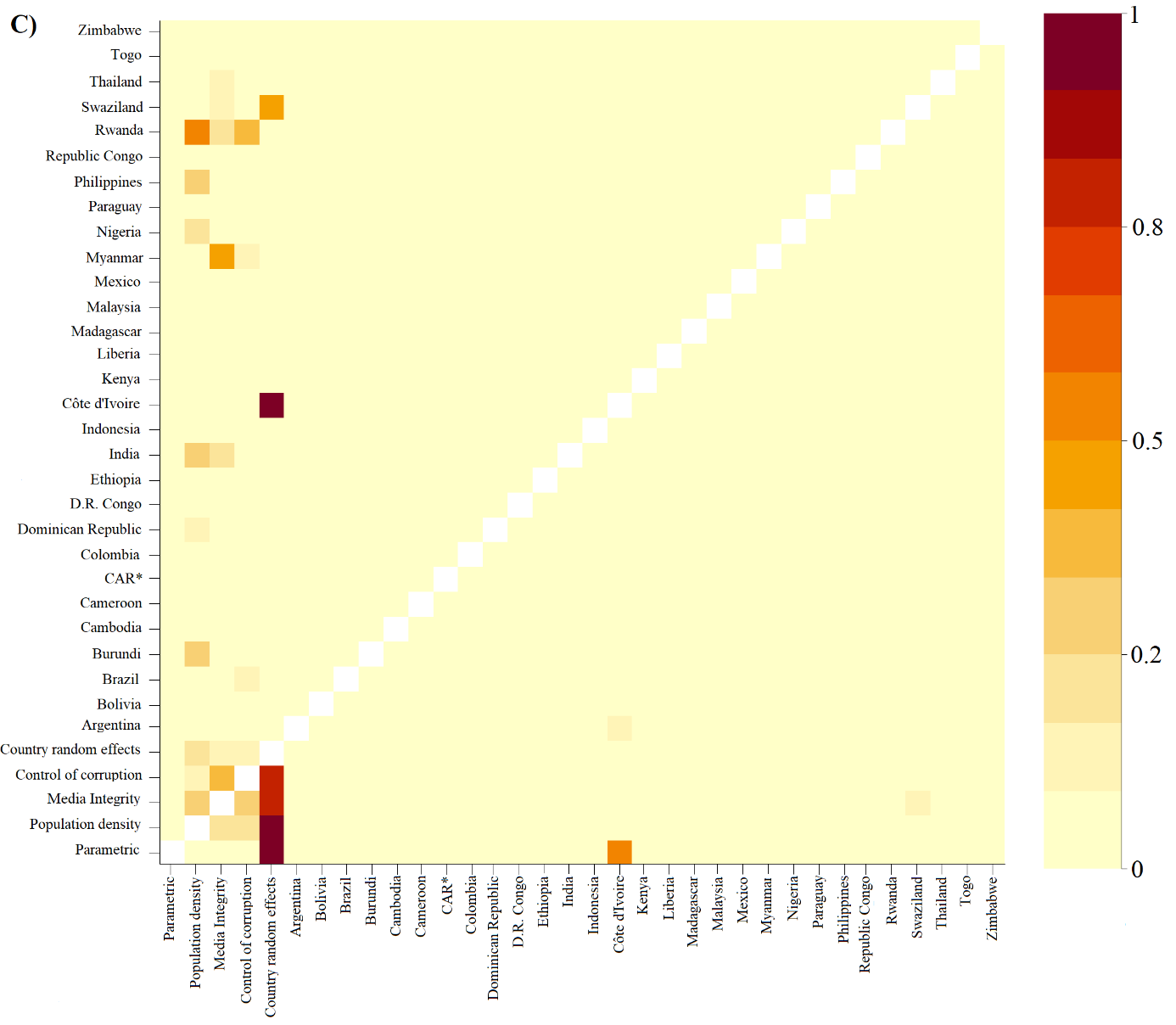

**Figure C3. Estimated concurvity from the Upper Chamber HGAM.** High values of concurvity were attained primarily in relation to the country random effects term. Population density, media integrity and control of corruption term have several moderate values of concurvity with multiple countries. The rest of the concurvity diagnostics have low values indicating that the model terms do not explain each other well. This suggest that the model does not have problems associated with high concurvity, which are similar to high co-linearity. * - Central African Republic.
