## Supplementary material for "The role of elections as drivers of tropical deforestation": Figure D1. Residuals

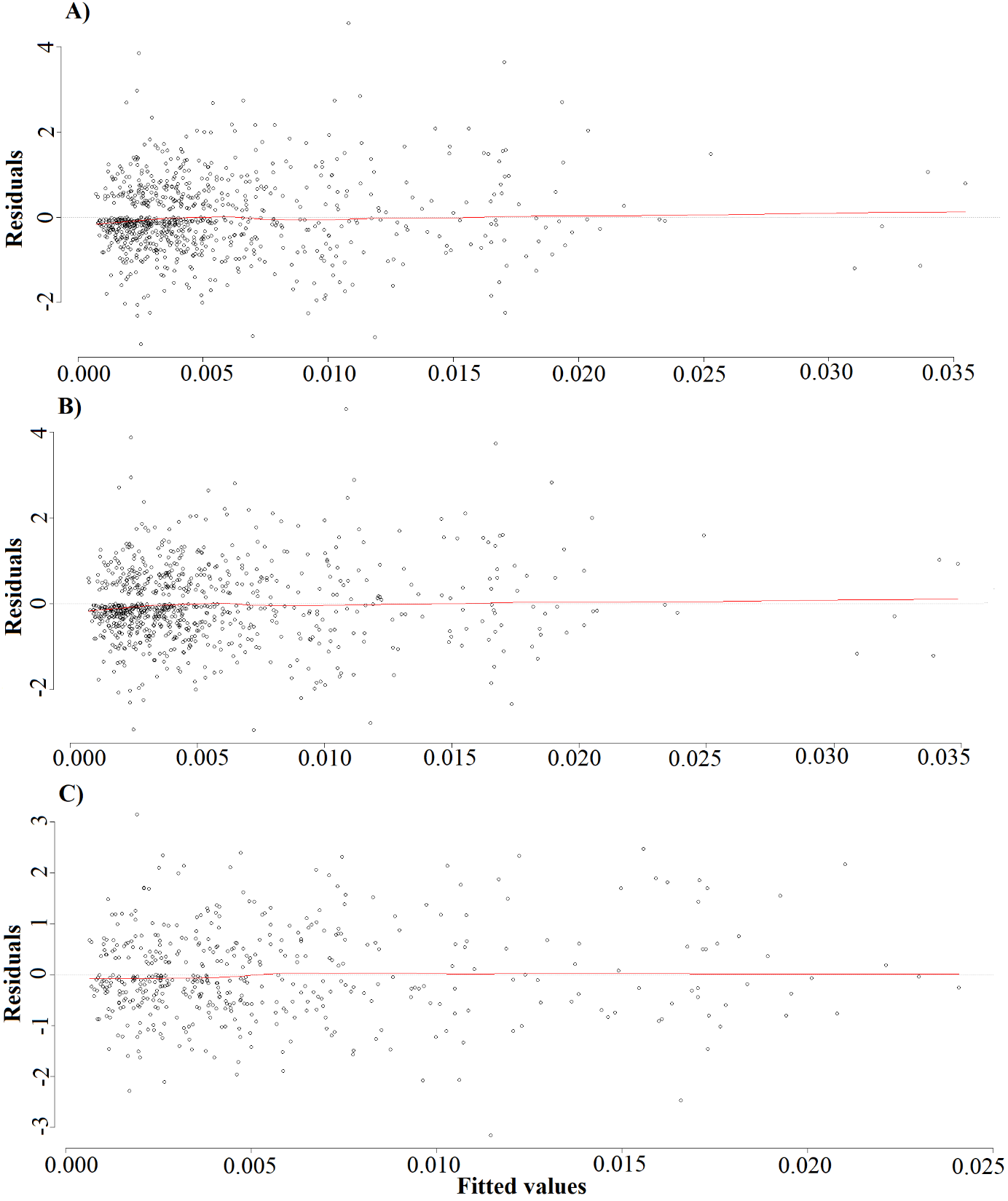

**Figure D1. Residual against the fitted values of the HGAMs.** The LOWESS line in red shows that the model fits the data quite well. The (A) presidential and (B) lower chamber HGAM have several extreme fitted values (>0.025) compared to the (C) upper chamber HGAM. The plots show no clear funnel or other trends, this indicates that the models fit well.
