## Supplementary material for "The role of elections as drivers of tropical deforestation": Table A1. Countries

**Table A1. Analysed countries and their respective deforestation from 2001–2019.** Countries in italics are excluded from the HGAM analysis.

| **Country** | **Deforestation (km^2^) *2001–2019*** |
| --- | --- |
| Angola  Argentina  *Belize*  Benin  Bolivia  Brazil  Burundi  Cambodia  Cameroon  Central African Republic  Colombia  Costa Rica  Dominican Republic  Democratic Republic Congo  Ecuador  Ethiopia  Fiji  *Gabon*  Ghana  Guatemala  Guinea  Guyana  *Honduras*  India  Indonesia  Ivory Coast  Kenya  Laos  Liberia  Madagascar  Malawi  Malaysia  Mali  Mexico  Myanmar  Nicaragua  Nigeria  Panama  Paraguay  Peru  Philippines  Republic Congo  Rwanda  Sierra Leone  Solomon Islands  *Suriname*  Swaziland  Tanzania  Thailand  Togo  Uganda  Venezuela  Vietnam  Zambia  Zimbabwe | 17531.6  37115.1  *2098.9*  995.1  41709.6  469839.0  162.6  17836.2  8866.7  5165.7  37635.8  2061.7  2534.9  112624.5  7280.2  2638.5  334.6  *3459.1*  6750.7  12328.2  7823.7  1763.2  *8937.0*  12700.3  227095.5  17094.1  2574.3  24172.6  10194.6  31825.8  1055.0  69899.9  450.7  30847.0  25703.5  13080.3  6995.1  3444.3  36877.1  27846.2  9693.7  5670.0  253.8  7768.5  1383.2  *1603.7*  728.0  15012.2  14885.2  405.9  5253.1  16795.3  20793.9  10234.3  1730.0 |
